# Module organizational principles and dynamics in biological networks

**DOI:** 10.1101/005025

**Authors:** Chun-Yu Lin, Tsai-ling Lee, Yi-Wei Lin, Yu-Shu Lo, Chih-Ta Lin, Jinn-Moon Yang

## Abstract

A module is a group of closely related proteins that act in concert to perform specific biological functions through protein–protein interactions (PPIs) that occur in time and space. However, the underlying organizational principles of a module remain unclear. In this study, we collected CORUM module templates to infer respective module families, including 58,041 homologous modules in 1,678 species, and PPI families using searches of complete genomic database. We then derived PPI evolution scores (PPIES) and interface evolution scores (IES) to infer module elements, including core and ring components. Functions of core components were highly correlated (Pearson’s *r* = 0.98) with those of 11,384 essential genes. In comparison with ring components, core proteins and PPIs were conserved in multiple species. Subsequently, protein dynamics and module dynamics of biological networks and functional diversities confirmed that core components form dynamic biological network hubs and play key roles in various biological functions. PPIES and IES can reflect module organization principles and protein/module dynamics in biological networks. On the basis of the analyses of gene essentiality, module dynamics, network topology, and gene co-expression, the module organizational principles can be described as follows: 1) a module consists of core and ring components; 2) the core components play major roles in biological functions and collaborate with ring components to perform certain functions in some cases; 3) the core components are conserved and essential in module dynamics in time and space.

## Introduction

Cooperation between proteins in time and space is essential for assembly of protein complexes that perform biological processes, such as cell cycle control and transcription (Gavin et al. 2006). Such protein assemblies can be regarded as a module, which often governs specific processes, such as natural variation, biological functions, and development, and is relatively autonomous with respect to other parts of the organism (Segal et al. 2003; Wagner et al. 2007). Global biological properties of modules have been analyzed in recent studies (Han et al. 2004; Gavin et al. 2006; Wagner et al. 2007; Kiel and Serrano 2012), and data from experimental methods (Gavin et al. 2006; Ruepp et al. 2008), network topology (Bader and Hogue 2003; Nepusz et al. 2012), gene expression-based methods (Ideker et al. 2002; Segal et al. 2003), and evolutionary analyses (Yamada et al. 2006) contribute to the concept of modularity. The proteins and protein-protein interactions (PPIs) in a module are often dynamics in time and space. They may change over seconds to assemble and disassemble the modules for cellular processes requirement, as well as evolve over millions of years as proteins and PPIs are gained and lost (Levy and Pereira-Leal 2008). Investigations of underlying organizational principles of protein modules are urgently required to improve the understanding of cellular processes and module evolution.

As complete genomes become increasingly available, systems biological approaches based on homologous PPIs and modules across multiple species may elucidate organizational principles, evolution, and dynamics of modules. An experimental genome-wide screen approach, based on the isoforms of complexes, was used to identify the 491 complexes, differentially combined with attachment proteins to execute time–space potential functions in yeast (Gavin et al. 2006). This work still limited in requirement of experimental methods and in one target organism (i.e., *Saccharomyces cerevisiae*). In addition, functionally interacting proteins have been shown to be gained or lost together during genome evolution (Ettema et al. 2001). However, functional modules showed limited conservation during evolution (Yu et al. 2004), with only 40% evolutionary cohesion among 1,161 Prokaryotic modules (Campillos et al. 2006). The causes of restricted evolutionary modularity are still unclear. Previously, we inferred the module family, which consists of a group of homologous modules, from complete genomic database (e.g. Integr8) through PPI families (Chen et al. 2007; Chen et al. 2009). Based on the module families and PPI families, we reconstructed module-module interaction networks (called MoNetFamily (Lin et al. 2012)) in vertebrates. However, the organizational principles and dynamics of modules in biological networks remain unclear.

To address these issues, we propose PPI evolution score (PPIES) and interface evolution score (IES) as the basis to study the organizational principles and characteristics of module in biological networks using module families and PPI families across multiple species. These two scoring systems reflect core and ring components of modules. Furthermore, we define protein dynamics (PDN) and module dynamics (MDN) of biological networks to reflect functional diversities of proteins and modules, respectively. According to PDN and MDN, proteins and PPIs of core components are often conserved in homologous modules and consistently play key temporal and spatial roles for performing biological functions. Conversely, ring proteins and PPIs are not often conserved in module families. Compared with ring proteins, core proteins are essential for survival, as indicated by the Database of Essential Genes (DEG) (Zhang and Lin 2009) and the Gene Ontology (GO) database (Ashburner et al. 2000), and preferentially constitute hubs of a PPI network. PPIES and IES values reflected evolutionary conservation and dynamics of modules and proteins in a human PPI network, which included 2,391 proteins and 11,181 PPIs. Moreover, core PPIs were co-expressed significantly more than ring PPIs in 7,208 *Homo sapiens* gene expression data sets (≥3 samples) from Gene Expression Omnibus (GEO) (Barrett et al. 2009). The present data indicate that core components of modules are temporal and spatial necessities for various biological functions, whereas ring components only participate in module functions in some cases. Finally, we suggest the following module organizational principles: 1) a module comprises core and ring components and the former is conserved across multiple species; 2) core components often play major roles in the biological functions of a module, whereas the ring components are indirectly involved in biological functions through collaborations with core components; 3) core components are often essential for temporal and spatial module dynamics, and for multiple cellular functions. We believe that our results are useful for understanding the module organizational principles in evolution, cellular functions, proteins, and module dynamics in biological networks.

## Results and Discussion

### Overview

Figure 1 shows the details of our method for identifying core and ring components of modules, and for elucidating module organizational principles through template-based homologous modules (module families) using the following steps (Fig. 1A): First, a module template database comprising 1,519 protein complexes (1,094 from *H. sapiens*, 248 from *Mus musculus*, 148 from *Rattus norvegicus*, and 29 from *Bos taurus*) was selected from the Comprehensive Resource of Mammalian protein complexes database (CORUM; release 2.0) (Ruepp et al. 2008). Internal PPIs of module templates were then added to templates that lacked PPIs using template-based homologous PPIs, including experimental PPIs from IntAct (Aranda et al. 2010), BioGRID (Stark et al. 2011), DIP (Xenarios et al. 2002), MIPS (Mewes et al. 2008), and MINT (Ceol et al. 2010), and predicted homologous PPIs (Chen et al. 2009; Lo et al. 2010) (Fig. 1B). For each PPI of a module, we inferred its PPI family with joint *E*-values of ≤10^−40^ (Chen et al. 2009) by searching a complete genomic database (Integr8 version 103, containing 6,352,363 protein sequences in 2,274 species) using previously identified homologous PPIs (Chen et al. 2009; Lo et al. 2010) (Fig. 1C). Subsequently, we utilized MoNetFamily (Lin et al. 2012) to identify homologous modules of module templates according to topological similarities across multiple species (Fig. 1D). Module profiles were then constructed for module families, and protein and PPI components were computed (Fig. 1E). Next, we then derived PPIES and IES scores to extrapolate core and ring components of a module. Finally, we constructed PPI networks and genome-wide investigations for organizational principles of a module, including gene essentiality (Fig. 1F), module dynamics (Fig. 1G), network topology (Fig. 1H), and gene expression profiles (Fig. 1I).

**Figure 1.**
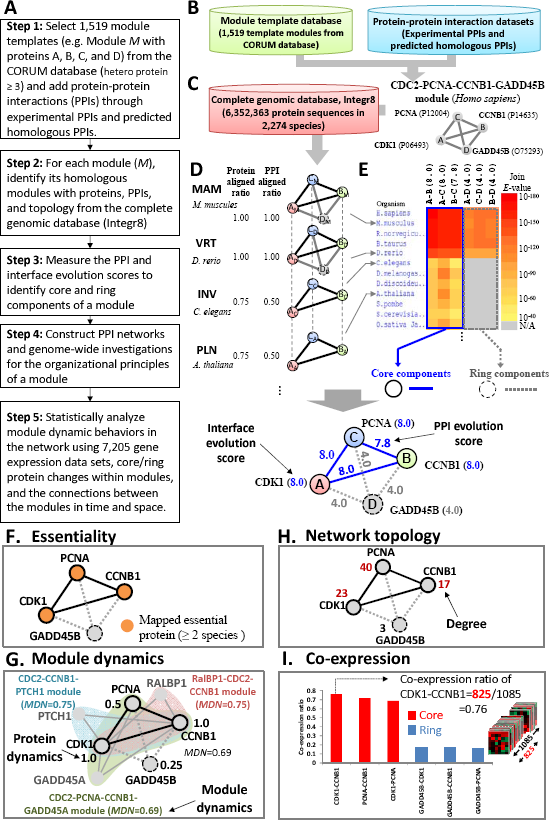
Overview of core and ring components of modules using the human CDC2–PCNA–CCNB1–GADD45B module as a template. (A) The main procedure. (B) Module template database and protein–protein interaction (PPI) database for inferring homologous PPIs. (C) Homologous PPIs and proteins of the template by searching the complete genomic database (Integr8). (D) Homologous modules of the CDC2–PCNA–CCNB1–GADD45B module. (E) PPI profiles and core (solid circle and line) and ring (dash circle and line) components of this module family across multiple organisms commonly used in molecular research projects. (F) Essentiality of core (solid circle) and ring (dash circle) proteins in this module family; orange circles indicate mapped essential proteins when they are homologs of ≥2 essential proteins. (G) The supermodule comprised four modules, including CDC2–PCNA–CCNB1–GADD45B, CDC2–CCNB1–PTCH1 (blue), CDC2–PCNA–CCNB1–GADD45A (green), and RalBP1–CDC2–CCNB1 (red), with their module dynamics. The dynamics of core proteins (e.g., CDK1 = 1.0) of this module are higher than those of ring proteins (i.e., GADD45B = 0.25). (H) The degrees of core and ring proteins of the module in the human PPI network, including 2,391 proteins and 11,181 PPIs. (I) Co-expressions of six PPIs (Pearson’s *r* ≥ 0.5) of this module were statistically derived from 7,208 microarray data sets.

### Homologous modules and human PPI network

To observe topologies and functional similarities of homologous modules in module families, we collected 75,706 homologous modules from 370 reference modules across 1,442 organisms from the KEGG MODULE database. According to the data set, protein-aligned ratios of 82% (62,080) between homologous and their reference modules were more than 0.5 (Supplemental Fig. S1A). To determine topological similarity thresholds between reference and its homologous modules, we added intra-module PPIs using the following PPI databases: 1) 461,077 experimental PPIs from annotated PPI databases, including IntAct (Aranda et al. 2010), BioGRID (Stark et al. 2011), DIP (Xenarios et al. 2002), MIPS (Mewes et al. 2008), and MINT (Ceol et al. 2010); 2) sequence-based homologous PPIs with joint *E*-values of ≤10^−40^ (Chen et al. 2009) among 461,077 experimental PPIs; and 3) 86,252 structure-based homologous PPIs with *Z*-scores of ≥4 (Chen et al. 2007). Among 75,706 organism-specific modules, we added at least one PPI for 23,092 modules, and 65% PPI-aligned ratios between reference modules and their homologous modules were ≥0.3 (Supplemental Fig. S1B). Here, we set the protein-aligned ratio and PPI-aligned ratio to 0.5 and 0.3, respectively, to identify homologous modules of a module template.

To derive homologous modules across multiple species, we collected 1,519 high-quality module templates, which are manually annotated protein complexes from the MIPS CORUM database. These 1,519 modules included 1,094 from *H. sapiens*, 248 from *M. musculus*, 148 from *R. norvegicus*, and 29 from *B. Taurus*, and contained at least three proteins (Ruepp et al. 2008). Based on these module templates and topology similarity thresholds of protein-aligned and PPI-aligned ratios, we inferred 58,041 homologous modules in 1,678 species from 461,077 sequence-based PPI families and 86,252 structure-based PPI families (Chen et al. 2007; Chen et al. 2009). Furthermore, we reconstructed the human PPI network using these 1,519 modules and their homologous modules, including 2,391 proteins and 11,181 PPIs.

### Core and ring components of a module

Homologous modules provide the clue to understand the evolution and conserved functions of proteins and PPIs within a module. Thus, we devised PPIES and IES of a module family to identify core and ring components. In a module family, a PPI with high PPIES indicate that its homologous PPIs are highly conserved across species and taxonomic divisions, such as mammals (MAM), vertebrates (VRT), invertebrates (INV), plants (PLN), bacteria (BCT), and archaea (ARC) according to the NCBI taxonomy database (see Methods). In addition, IES of the protein *i* was set to the maximum PPIES of these PPIs, which reflected interactions between protein *i* and its partners. Here, we considered proteins with IES ≥ 7 and PPIs with PPIES ≥ 7 as core components of the module, and other proteins and PPIs are referred to as ring components.

We used the CDC2–PCNA–CCNB1–GADD45B module family as an example to illustrate core and ring components and their biological properties (Figs. 1D and 1E). The core components of the CDC2–PCNA–CCNB1–GADD45B module (CORUM ID: 5545 [31]) included three proteins (solid circles; i.e., cyclin-dependent kinase 1 (CDK1/CDC2), proliferating cell nuclear antigen (PCNA), and G2/mitotic-specific cyclin-B1 (CCNB1)), with IESs of 8.0, and three PPIs (solid lines; i.e., CDK1–CCNB1 and CDK1–PCNA with PPIESs of 8.0, and CCNB1-PNCA with a PPIES of 7.8). Ring components (dashed circles and lines) consist of the growth arrest and DNA damage-inducible protein (GADD45) with an IES of 4.0 and three PPIs (GADD45–CDK1, GADD45–PCNA, and GADD45–CCNB1) with PPIESs of 4.0. During the G2/M cell cycle phase, GADD45B specifically interacts with the CDK1–CCNB1 complex, but not with other CDK–Cyclin complexes, to regulate activation of G2/M cell cycle checkpoints (Vairapandi et al. 2002).

According to PPI profiles of this module in organisms that are commonly used in molecular research projects (Fig. 1E), we found that core PPI families of CDK1–CCNB1, CKD1–PCNA, and CCNB1–PNCA were highly conserved. For example, the profile of the CDK1–CCNB1 PPI is conserved across 67 species according to the interaction between CDK1 and CCNB1 of *H. sapiens* that were shown purification by protein kinase assays (Aranda et al. 2010) and co-immunoprecipitation experiments (Xenarios et al. 2002). During the G2 cell cycle phase, active CDK1–CCNB1 interactions enhance chromosome condensation, and nuclear envelope breakdown into separate centrosomes (Lindqvist et al. 2007). During the response to DNA damage, PCNA is conveniently positioned at the replication fork to coordinate DNA replication, and activates DNA repair and damage tolerance pathways. However, no GADD45B (ring protein) homologs were found in chloroplasts or bacteria. GADD45B is involved in G2/M cell cycle arrest, acting as an inhibitor of the CDK1–CCNB1 complex, and contributes to regulation of S and G2/M cell cycle checkpoints following exposure of cells to genotoxic stress (Vairapandi et al. 2002). These results indicate that core components often play major roles in temporal and spatial biological functions of the module, whereas ring components may collaborate with core components to contribute to certain functions.

### Essential proteins and protein interface evolution scores

Essential genes (or proteins) are considered to be required to support cellular life and likely to be common to all cells (Kobayashi et al. 2003). To evaluate essentiality of core and ring proteins in module families, we collected 11,384 essential proteins over 25 species from DEG (version 6.5) (Zhang and Lin 2009), including 8 eukaryotes (e.g. *H. sapiens* and *S. cerevisiae*) and 17 prokaryotes (e.g. *Escherichia coli* and *Bacillus subtilis*). Because homologs of essential proteins are likely to be essential, module proteins were considered essential when they were homologous to those recorded in DEG. For example, CCNB1 is a mapped essential protein and is homologous to essential proteins B_M_ (G2/mitotic-specific cyclin-B1 in mouse) and B_D_ (cyclin B1 in zebrafish) from DEG (Figs. 1D and 1F). For the CDC2–PCNA–CCNB1–GADD45B module family, homologs of the core proteins CDK1, CCNB1, and PCNA were essential proteins according to DEG (Zhang and Lin 2009). In contrast, all homologs of the ring protein GADD45B were non-essential (Fig. 1F).

According to the DEG data set, 7,950 proteins from 1,519 module templates were clustered into two groups, including 3,628 mapped essential proteins and 4,322 unannotated proteins that were absent from DEG. Among these 3,628 mapped essential proteins, IES values of 60% are more than 7 and their *IES* values are significantly higher than those of unannotated proteins (Mann–Whitney U test, P = 3e-217; Fig. 2A). In addition, percentages of mapped essential proteins were significantly correlated with IES (Pearson’s *r* = 0.98) and these increased rapidly with IES ≥ 7 (Fig. 2B). Interestingly, IES values of 968 mapped essential proteins that were conserved over two species (“mapped ≥ 2 species”) were much higher than those of the 3,628 mapped essential proteins that were conserved over only one species (“mapped ≥ 1 species”; P = 2e-72; Supplemental Fig. S2).

**Figure 2.**
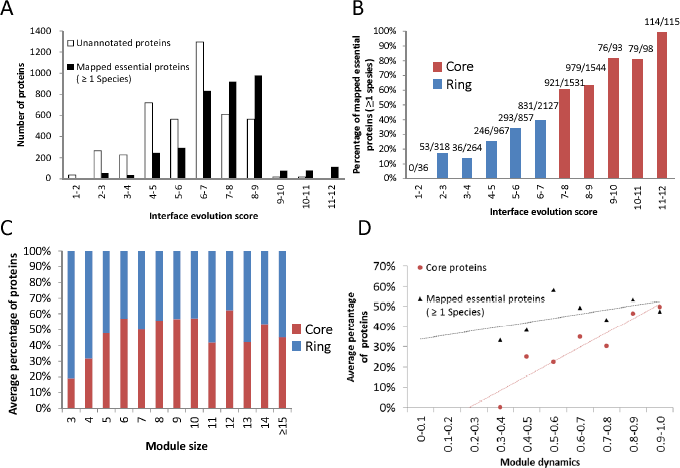
Characteristics of module organizational principles and dynamics. (A) Interface evolution score (IES) distributions of the numbers of unannotated (white) and mapped essential proteins (≥1 species, black). (B) The relationship between IES values and percentages of mapped essential proteins (≥1 species), showing significant increases when the IES is ≥7 (red). In this study, proteins with IES values of ≥7 and <7 were considered core proteins (red) and ring proteins (blue), of a module, respectively. (C) The relationship between module sizes and core/ring composition of modules; percentages of core and ring components in different module sizes are similar. (D) Pearson’s *r* values between module dynamics and percentages of core proteins (red) and mapped essential proteins (black) were 0.93 and 0.52, respectively.

Based on these 11,384 essential proteins, we derived 160 essential GO molecular function (MF) terms (Supplemental Table S1, Supplemental Figs. S3 and S4, and Supplemental Text 1) and analyzed functional annotations of core and ring components using hypergeometric distributions (P ≤ 0.05). Among 160 essential GO MF terms, 31% were involved in central dogma (Supplemental Fig. S3A), and 21% (33 terms, such as acetyl-CoA carboxylase activity) were recorded for carbohydrate and lipid metabolism. In addition, 16 terms were involved in amino acid metabolism (e.g., cysteine desulfurase activity) and RNA degradation (e.g., 3′–5′ exonuclease activity). Whereas the distribution of occurrence ratios of these 160 terms did not differ between the core component set and the essential protein set (Pearson’s *r* = 0.77), and that between the ring component set and the essential protein set differed significantly (Pearson’s *r* = 0.49; Supplemental Fig. S4). Specifically, both core and essential protein sets had some significant MF terms, such as “structural constituent of ribosome,” “ATPase activity,” “nucleoside-triphosphatase activity,” and “chromatin binding.” These terms commonly relate to processes that are critical for survival and are conserved in the modules.

Using orthologs from the PORC database (Kersey et al. 2005) and these 160 essential GO MF terms, we analyzed 1,212 unannotated core proteins (IES ≥ 7; Table 1). Among these, 462 (38%) were orthologous to essential proteins or were annotated with at least one of the 160 essential GO MF terms. Furthermore, 303 unannotated core proteins (25%) possessed child annotations of the 160 essential GO MF terms; therefore, were considered essential. Moreover, 76% and 100% of the unannotated core proteins with IES ≥ 9 or 11, respectively, were annotated with orthologs of essential proteins, were one of 160 essential GO MF terms, or were child annotations of the 160 essential GO MF terms (Table 1). These results show that protein IES provides biological insights, and that core components are often essential for survival, as indicated in DEG (Zhang and Lin 2009) and GO (Ashburner et al. 2000).

**Table 1.** Summary of interface evolutions, orthology, and essential molecular functions

| Interface evolution score | Number of total proteins | Number of annotated proteins (recorded as essential genes) | Number of unannotated proteins | Orthologs (PORC)<sup>a</sup> | 181 essential GO MF terms<sup>b</sup> | Children of 181 essential GO MF terms<sup>c</sup> | Total |
| --- | --- | --- | --- | --- | --- | --- | --- |
| ≥ 11 | 115 | 114<br>(99%) | 1 | 0 | 0 | 1 | 1<br>(100%) |
| ≥ 9 | 306 | 269<br>(88%) | 37 | 14 | 10 | 19 | 28<br>(76%) |
| ≥ 7 | 3381 | 2169<br>(64%) | 1212 | 152 | 368 | 303 | 599<br>(49%) |
| < 7 | 4569 | 1459<br>(32%) | 3110 | 217 | 491 | 689 | 1175<br>(38%) |

<sup>a</sup> The number of proteins contained at least an orthologous protein, recorded in the PORC orthology database (Kersey et al. 2005), of essential proteins.
<sup>b</sup> The number of proteins contained at least an essential GO MF terms.
<sup>c</sup> The number of proteins contained at least a child term of 181 essential GO MF terms..

Figure 2C shows the relationship between module sizes and core/ring compositions of modules. In a module, the number of core components is similar (∼50%) to the number of ring components when the module size ≥5. We next analyzed the distributions of three kinds of modules: including core-only module, ring-only module, and core-ring module. Interestingly, the percentages of core-only modules were often less than 18% and were much lower than those of ring-only modules (Supplemental Fig. S5). In previous studies, functional modules had limited conservation during evolution (Yu et al. 2004), with approximately 40% of 1,161 prokaryotic modules displaying evolutionarily cohesion (Campillos et al. 2006). The present analyses suggest that this limitation of evolutionary modularity may reflect low percentages of core-only modules and prevalence of core–ring modules (approximately 50%). In addition, we found that small modules (approximately 71%) prefer to contain only ring components (ring-only modules). This observation suggests that the size of young modules is often smaller than that of ancient modules (Campillos et al. 2006).

### Protein dynamics and module dynamics in a network

In the present study, supermodules comprised several modules, often with specific biological functions, and their functional diversity was defined by numbers of modules. Initially, 1,515 human CORUM modules were clustered into supermodules using the Jaccard similarity coefficient *J*(A,B) (Willet 1998). The *J*(A,B) is defined as 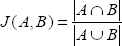, where *A* ∩ *B* is the number of common proteins (intersection set) in modules A and B, and *A* ∪ *B* is the number of the union protein set in modules A and B. Here, modules A and B are clustered into one group if *J*(A,B) ≥ 0.5. Based on this threshold, we iteratively clustered modules and groups into supermodules until *J*(A,B) ≤ 0.5 for any pair of modules (or groups). Finally, we clustered 1,515 modules into 185 supermodules (including 736 modules) and 85 supermodules (including 338 modules) when the numbers of modules in a supermodule are more than 2 and 3 modules, respectively. Specifically, the CDC2–PCNA–CCNB1–GADD45B module was grouped with 3 other experimental modules to form the CDC2–PCNA–CCNB1 supermodule, which included RalBP1–CDC2–CCNB1, CDC2–CCNB1–PTCH1, and CDC2–PCNA–CCNB1–GADD45A modules (Fig. 1G). The functional diversity of the CDC2–PCNA–CCNB1 supermodule was 4.

Proteins often assemble dynamically and cooperate to form the modules that perform biological functions in time and space. Among 1,515 human modules, we found that 1,449 (96%) contained at least one protein that was involved in more than two modules. Therefore, it is assumed that the importance of a protein is reflected by the number of modules that it is involved in. To assess dynamics (PDN) of a core and ring protein in a supermodule (or in a biological network), we defined 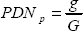 of the protein *p*, where *g* is the number of modules in which protein *p* is involved and G is the number of modules in the supermodule. Thus, a protein *p* that is involved in all biological functions of a supermodule has a *PDN_p_* of 1. Subsequently, mean PDN of modules were calculated as a measure of MDN, and module *m* in a biological network is given as 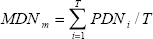, where *T* is the number of proteins in the module *m*. High MDN implies that the module highly participates in various functions in the biological network and often plays an important role in a cell.

Figure 2D shows the correlation of MDN values with percentages of core proteins (Pearson’s *r* = 0.93) and mapped essential proteins (Pearson’s *r* = 0.52) in modules. Among 185 supermodules with ≥2 modules (736 modules; Fig. 3A), three of four proteins (75%) in the CDC2–PCNA–CCNB1–GADD45B module, which has a high MDN value (0.69) in its supermodule, were both core proteins and mapped essential proteins (Fig. 1G). Similarly, the correlation between MDN and average module evolution scores (MES) was significant (Pearson’s *r* = 0.91; Fig. 3B). In the CDC2–PCNA–CCNB1 supermodule, MDN values of four modules were ≥0.69 and represented high MES (≥6). These results indicate that modules that participate in various functions in a biological network are often essential and conserved.

**Figure 3.**
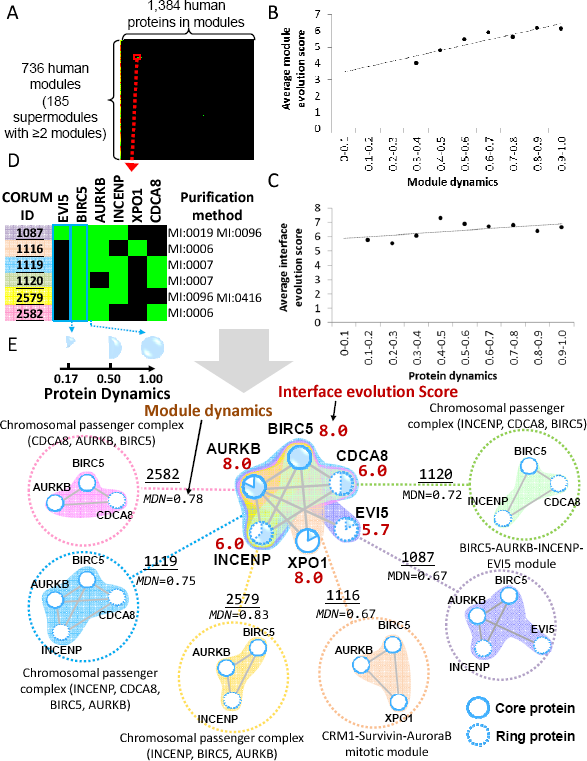
Module groups and the chromosomal passenger complex (CPC) supermodule. (A) Clustering matrix of 736 human modules and 1,384 proteins. The distribution of (B) module dynamics (MDN) and (C) protein dynamics (PDN) against interface evolution scores (IES) and module evolution scores (MES), respectively. MDN were highly correlated (Pearson’s *r* = 0.91) with average MES. Similarly, PDN and average IES values were significantly correlated (Pearson’s *r* = 0.56). (D) The profile of the CPC supermodule with six experimental modules that were identified previously using various purification methods; (E) the CPC supermodule comprises six CORUM modules; MDN, PDN, and IES (red) are shown. Solid and dashed circles denote core proteins and ring proteins, respectively.

In addition, PDN were correlated with average protein IES values (Pearson’s *r* = 0.56; Fig. 3C), and the dynamics of core proteins (IES ≥ 7) were significantly higher (Mann–Whitney U test, P = 5e-10) than those of ring proteins (IES < 7). Among 85 module groups comprising ≥3 modules, the average protein dynamic values of core and ring proteins were 0.70 and 0.56, respectively. In the CDC2–PCNA–CCNB1 supermodule, core proteins were involved in multiple modules (PDN ≥ 0.5), whereas ring proteins were not (PDN = 0.25; Fig. 1G). These results suggest that core components play major roles in biological functions in time and space, and that ring components sometimes participate in module functions.

The chromosomal passenger complex (CPC) supermodule comprises six experimental modules that were derived from various purification methods, including anti bait coimmunoprecipitation (MI:0006), anti tag coimmunoprecipitation (MI:0007), coimmunoprecipitation (MI:0019), pull down (MI:0096), and fluorescence microscopy (MI:0416) (Fig. 3D). This supermodule is organized by six proteins, including aurora-B serine/threonine protein kinase (AURKB), baculoviral IAP repeat-containing protein 5 (BIRC5; Survivin), inner centromere protein (INCENP), borealin (CDCA8), ecotropic viral integration site 5 protein homolog (EVI5), and exportin-1 (XPO1/CRM1; Figs. 3D and 3E). During early mitosis, CPC is an important mitotic regulatory complex that promotes chromosome alignment by correcting misattachments between chromosomes and microtubules of the mitotic spindle (Vader et al. 2006b). The CPC supermodule contained the three core proteins BIRC5, AURKB, and XPO1, and the three ring proteins INCENP, CDCA8, and EVI5. In this supermodule, chromosomal passenger complex (INCEP, AURKB, and BIRC5) had the highest MDN value (0.83), and comprised two core proteins and three essential proteins.

Interestingly, the MDN value of the CRM1–Survivin–AuroraB mitotic module (BIRC5, AURKB, and XPO1) was 0.67, and its module evolution score was 8. The core proteins BIRC5 and AURKB were included in most CPC modules (PDN ≥ 0.83), whereas PDN of XPO1 was only 0.17 (Fig. 3E). The functions of the CPC can attribute to the action of the enzymatic core, the AURKB (Vader et al. 2006b), and the BIRC5 mediates the CPC to target to the centromere and midbody (Vader et al. 2006a). Previous studies indicate that the BIRC5–XPO1 interaction is essential for CPC localization and activity (Knauer et al. 2006), implying that XPO1 may play an important role.

PDN values of the ring proteins INCENP, CDCA8, and EVI5, were 0.67, 0.5, and 0.17, respectively (Fig. 3E). In human cells, functional CPCs can be targeted, although less efficiently, to centromeres and central spindles in the absence of CDCA8, lack of orthologs in *S. cerevisiae* and *S. pombe*, when BIRC5 is linked covalently to INCENP (Vader et al. 2006a). During the late stages of mitosis, EVI5 associates with CPC and plays a role in the completion of cytokinesis. Therefore, the present results suggest that the dynamics of core proteins are often significantly higher than those of ring proteins.

### Network topology and protein interface evolution scores

To analyze core and ring components in PPI networks, we derived a human PPI network from 1,515 homologous modules. This PPI network comprised 2,391 proteins and 11,181 PPIs (Figs. 4A and 4B), and is a scale-free network that can be described as *P*(*k*) ∼ *k^−r^*, in which the probability of a node with *k* links decreases as the node degree increases on a log–log plot (Fig. 4C). The degree exponent γ was 1.60 in this PPI network, which was consistent with the architecture (weak scale-free network properties) of previously described cellular networks (Barabasi and Oltvai 2004; Seyed-allaei et al. 2006). Figure 4C shows the distribution of node degrees for core proteins, ring proteins, and all proteins in this human PPI network. For 1,069 core proteins, 1,322 ring proteins, and 2,391 proteins of this PPI network, the distribution of node degrees of core proteins (mean, 15.12) was significantly higher than that of ring proteins (mean, 4.69), and that of all proteins (mean, 9.35). Moreover, the median and top 25% of degrees of this network were 4 and 10, respectively.

**Figure 4.**
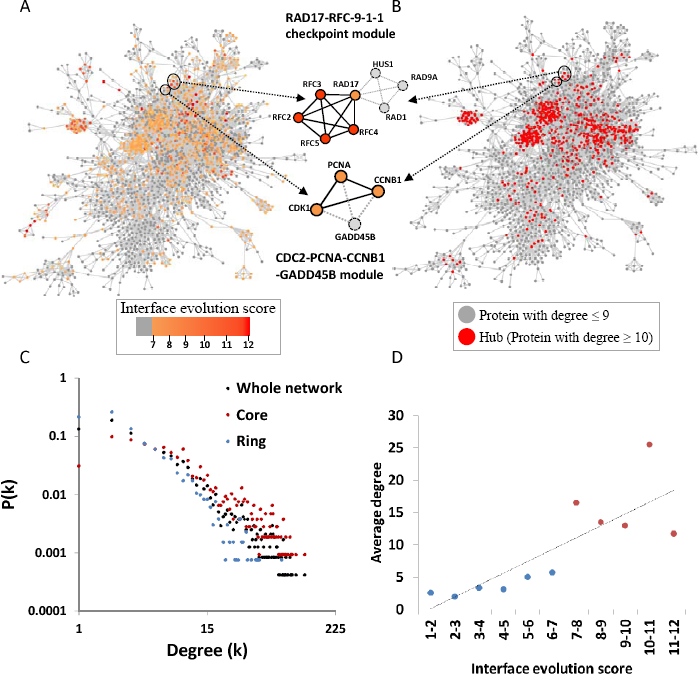
Topologies of core and ring proteins in the human protein–protein interaction network comprising 2,391 proteins and 11,181 PPIs. Each protein of the network is annotated with (A) interface evolution scores (IES) and (B) degrees. The core proteins, such as RFC2, RFC3, RFC4, and RFC5, are often hubs (top 25% of the highest degree) of this network. (C) Node degree distributions of all proteins (black), core proteins (red), and ring proteins (blue) in this scale-free PPI network. (D) The distribution of IES values against average degrees of proteins; IES values of proteins were highly correlated (Pearson’s *r* = 0.81) with average node degrees. Core proteins (red) have higher degrees than ring proteins (blue), and constitute the hubs of PPI networks.

On the basis of a previous study (D’Antonio and Ciccarelli 2011), we considered proteins within the top 25% of the highest degree (here, degree ≥10) as hubs of the network. The IES distribution of these core proteins was consistent with the hub distribution of this PPI network, particularly at the center of the network (Figs. 4A and 4B). Moreover, 43% of core proteins with degrees of ≥10 were hubs, and only 12% of ring proteins were hubs. Protein IES values were also highly correlated (Pearson’s *r* = 0.81) with average node degrees (Fig. 4D). Interestingly, node degrees of ring proteins in modules were lower than those of all proteins in this network, indicating that core proteins but not ring proteins play major roles in high connectivity of module sub-networks. Our results suggest that core proteins are preferential constituents of network hubs, as reflected by protein IES values. This observation is consistent with a previous study showing that highly conserved enzymes in a metabolic network were frequently highly connected at the center of the network and were involved in multiple pathways (Peregrin-Alvarez et al. 2009).

In the CDC2–PCNA–CCNB1–GADD45B module, the core proteins CDK1, CCNB1, and PCNA had higher degrees (≥17) than the ring protein GADD45B (degree = 3) in the human PPI network (Figs. 1H, 4A, and 4B). Therefore, the core components of module families preferentially comprise hubs of the PPI network and are essential elements for the survival of an organism.

### Connectivity of modules

A module is relatively autonomous and often has high connectivity (*C_t_*) within a PPI network. To observe connectivity (*C_t_*) of a module in a PPI network, we quantified the connectivity by 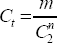 (Campillos et al. 2006), where *n* and *m* are the numbers of connected proteins and PPIs in a module. A *C_t_* value of 1 indicates that proteins are completely interconnected in a module. Here, we computed *C_t_* of modules using the human PPI network. Figure 5A shows the *C_t_* of core and ring components, module templates, and their respective extended modules (four types of modules). Here, modules were extended by one-layer of PPIs and proteins in the module template. Among these four module types, *C_t_* values of core compounds was significantly higher than the others, and that of extended modules was the lowest. Among 1,519 module templates, *C_t_* values of more than 0.6 were observed in 71% (1,081) of cases. In contrast, *C_t_* values were more than 0.6 for only 5% (71) of extended modules. Moreover, 90% of core components and 81% of ring components had *C_t_* values of ≥0.6. Similarly, 58,041 modules that were homologous to module templates had *C_t_* values of ≥0.6 in 76% of cases (44,319), whereas only 1% (842) of their extended modules had *C_t_* values of ≥0.6 (Supplemental Fig. S6A). These results indicate that core components have the highest connectivity, and that the modules in this PPI network have high connectivity.

**Figure 5.**
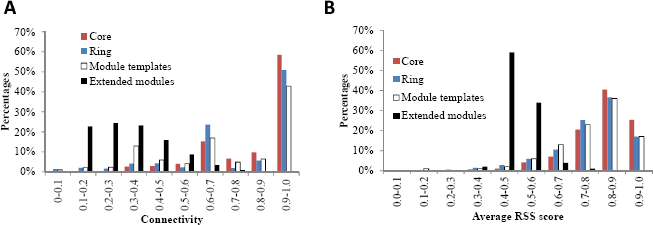
Distributions of connectivity (*C_t_*) and average relative specificity similarity (RSS) of modules in a human protein–protein interaction network. (A) *C_t_* distributions of core and ring components, module templates, and extended modules using 1,519 module templates; the extended module is a sub-network that includes one-layer extensions of PPIs and proteins of the module template. (B) Distributions of average RSS scores of GO biological processes for core and ring components, module templates, and extended modules. For *C_t_* values and average RSS, the core component (red) is the highest, and the extended module is the lowest (black); templates (white) are much higher than their extended modules.

### Biological functions of modules

Through assembly and cooperation of proteins in a PPI network, components of a module simultaneously perform certain biological functions. Based on the relative specificity similarity (RSS) (Wu et al. 2006) of GO terms, such as biological process (BP) and cellular component (CC), we computed AvgRSS scores to assess shared biological functions of all protein pairs in a module. The AvgRSS is defined as 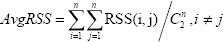, where *i* and *j* are any two proteins of a module and *n* is the number of proteins in the module.

To elucidate biological functions of modules, we compared module templates, their core and ring components, and their extended modules. For 1,519 module templates, BP and CC AvgRSS scores were more than 0.6 in 89% and 97% of cases, respectively (Fig. 5B and Supplemental Fig. S6C), and these scores were significantly higher than those of extended modules. In addition, BP and CC AvgRSS scores of core components were the highest among these four types. CC AvgRSS scores (97%) of templates were slightly higher than those of their ring components (94%) with AvgRSS scores of ≥0.6. Furthermore, BP and CC AvgRSS scores were more than 0.6 for 81% and 94% of homologous modules, respectively (Supplemental Figs. S6B and S6D). Similarly, BP and CC AvgRSS scores for core components of homologous modules were also significantly higher than those of ring components. Specifically, BP and CC AvgRSS scores for the CDC2–PCNA–CCNB1–GADD45B homologous module in *H. sapiens* were 0.79 and 0.84, but for extended modules they were only 0.43 and 0.25, respectively. The core components of this module had high BP and CC AvgRSS scores of 0.89 and 0.85, respectively. These results indicate that homologous modules of a template have highly similar biological functions and that their core components regulate similar biological processes and are often localized to the same cellular compartments.

### Co-expression behavior of core and ring components

Dynamic assembly and cooperation of proteins in time and space is essential for biological processes in a cell. In this study, we found that modules can be organized into core and ring components, which represent temporal and spatial conservation of dynamic PPIs and proteins. Genome-wide gene expression profiles are descriptive of molecular states that are associated with various responses to environmental perturbations and cellular phenotypes (Carter et al. 2004). Thus, to observe the dynamics of PPIs and proteins in a module, we collected 7,208 *H. sapiens* gene expression data sets (≥3 samples) from GEO (Barrett et al. 2009). Figures 6A and 6B show the details of the procedures for preparation and normalization of these data sets. For each module among 1,519 temples, we initially selected gene expression sets that contain all proteins in this module, and evaluated co-expressions of intra-module PPIs to construct a correlation matrix (Fig. 6C). To confirm that modules in the data sets were associated with biological functions, we selected gene expression sets that give rise to comparatively high protein expression and contain at least one co-expression of intra-module PPIs with Pearson’s *r* values of ≥*h* (see Methods).

**Figure 6.**
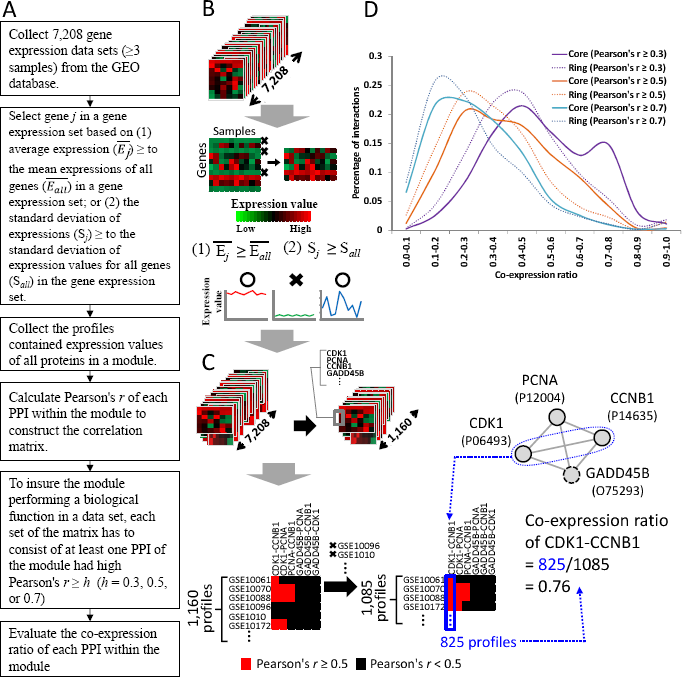
Gene co-expressions of core and ring PPIs in the modules using 7,208 *H. sapiens* gene sets from Gene Expression Omnibus (GEO). (A) The main procedure for collecting gene profiles and evaluating co-expression of core and ring PPIs in modules. (B) Gene expression profiles are collected by discarding non-significant genes with low expression and low expression variance. (C) Co-expression profiles of all protein pairs (PPIs) from the CDC2–PCNA–CCNB1–GADD45B module. (D) Distributions of co-expressions of core PPIs (solid lines) and ring PPIs (dot lines) of 1,515 human modules based on Pearson’s *r* thresholds of ≥0.3, ≥0.5 and ≥0.7. Co-expressions of core protein pairs are significantly higher than those of ring protein pairs.

Figure 6D shows relationships between co-expression ratios (CE) with Pearson’s *r* values of ≥0.3, 0.5, and 0.7 and percentages of core PPIs and ring PPIs for 1,515 human modules. When Pearson’s *r* values were ≥0.3, the average CE (0.51) of interacting core proteins (core PPIs) was significantly higher than that (0.44) of interacting ring proteins (ring PPIs; Mann–Whitney U test, P = 3e-79). Similarly, when Pearson’s *r* values were ≥0.7, the CE of interacting core proteins remained significantly higher than the ratio of interacting ring proteins (P = 3e-14). Specifically, the core PPIs CDK1–PCNA, PCNA–CCNB1, and CDK1–CCNB1 in the CDC2–PCNA–CCNB1–GADD45B module had significantly higher CEs (≥0.69) than those of the ring PPIs (≤0.18) CDK1–GADD45B, CCNB1–GADD45B, and PCNA–GADD45B, according to 1,085 high expression profile sets for this module (Fig. 1I). These results indicate that core PPIs of modules are co-expressed more frequently than ring PPIs, suggesting that core components are often simultaneously active or inactive in time and space.

### RAD17–RFC-9-1-1 checkpoint module

In this study, we used the RAD17–RFC-9-1-1 checkpoint module (RAD17–RFC-9-1-1 module, CORUM ID: 274) of *H. sapiens* to describe module organizational principles, protein dynamics, and module dynamics in biological networks. This module comprises 16 PPIs and 8 proteins (Fig. 7A), including the cell cycle checkpoint proteins RAD1/RAD9A/RAD17 (RAD1/RAD9A/RAD17), the replication factor C subunits 2/3/4/5 (RFC2/RFC3/RFC4/RFC5), and checkpoint protein HUS1 (HUS1). During the cell cycle, the RAD17–RFC-9-1-1 module is involved in the early steps of the DNA damage checkpoint response (Bermudez et al. 2003). Using the RAD17–RFC-9-1-1 module in *H. sapiens* as a module template, homologous modules across 127 species and 5 taxonomic divisions were all found to regulate DNA damage recognition (Fig. 7B). The ten PPI families (e.g., RFC2–RFC5, RAD17–RFC4, and RFC3–RFC4) and the six PPI families (e.g., HUS1–RAD9A and HUS1–RAD1) of this module were regarded as core components and ring components (Fig. 7C), respectively.

**Figure 7.**
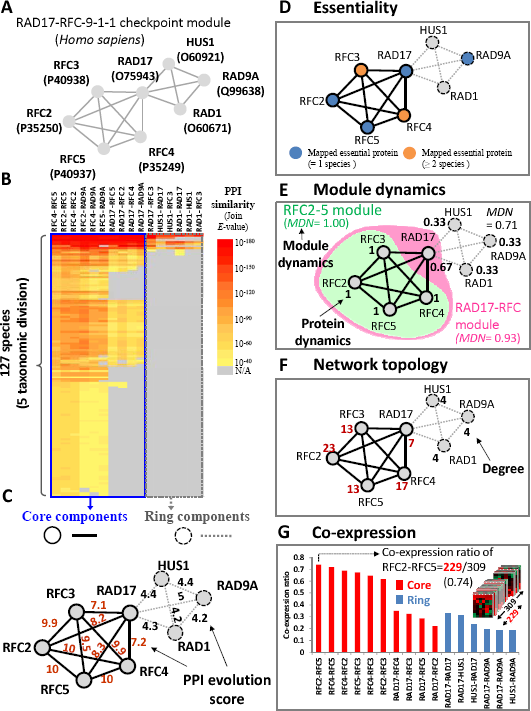
Characteristics and dynamics of core and ring components of RAD17–RFC-9-1-1 checkpoint module. (A) The RAD17–RFC-9-1-1 checkpoint module. (B) The module family profile includes 8 proteins and 16 PPI families. (C) Solid circles and lines denote the 5 core proteins and 10 core PPIs, respectively, and dashed circles and lines denote the 3 ring proteins and 6 ring PPIs, respectively. The PPI evolution scores are indicated. (D) Blue and orange circles indicate essential proteins when ≥1 and ≥2 mapped essential proteins are present, respectively. (E) The RAD17–RFC-9-1-1 supermodule comprises three modules with protein/module dynamics, including RFC2–5 (green), RAD17–RFC (pink), and RAD17–RFC-9-1-1. (F) Degrees of core and ring proteins in the human PPI network, including 2,391 proteins and 11,181 PPIs. (G) Co-expression ratios of 16 PPIs among 309 expression profiles selected from 7,208 gene sets. The solid circle and line denote the core protein and PPI, respectively. The dash circle and line indicate the ring protein and PPI, respectively.

Five core proteins, RFC2, RFC3, RFC4, RFC5, and RAD17, were homologous to essential proteins recorded in DEG (Fig. 7D) and annotated with several essential GO MF terms, such as “DNA clamp loader activity” and “nucleoside-triphosphatase activity.” During DNA replication, RFC binds to primed templates and recruits PCNA to the site of replication (Waga and Stillman 1998). In addition, RAD17 associates with these four small RFC subunits and forms an RFC-like complex that acts as a DNA damage sensor (Bermudez et al. 2003). Therefore, the present results suggest that core proteins of RFC subunits and RAD17 are essential in the RAD17–RFC-9-1-1 module.

The RAD17–RFC-9-1-1 supermodule comprises the RFC2–5 module (CORUM ID: 2200), the RAD17–RFC module (CORUM ID: 270), and the RAD17–RFC-9-1-1 module. Dynamic values (PDN = 1) of the core proteins RFC2, RFC3, RFC4, and RFC5 were consistently involved in these 3 modules to perform various biological functions (Fig. 7E). Conversely, the PDN value of the three ring proteins was 0.33, and these are included in one module to perform one of functions of the RAD17-RFC-9-1-1 supermodule. Moreover, module dynamic values of RAD17–RFC-9-1-1, RAD17–RFC, and RFC2–5 modules were 0.71, 0.93, and 1.0, respectively, and were highly correlated with MES (7.32, 8.99, and 9.81, respectively). In addition, the core proteins RFC2 (degree = 23), RFC3 (degree = 13), RFC4 (degree = 17), and RFC5 (degree = 13) were determined as hubs (degree ≥ 10) in the human PPI network (Fig. 7F). Conversely, the degree of all ring proteins (HUS1, RAD1, and RAD9A) was 4.

Among collected 7,208 gene expression data sets of *H. sapiens*, 309 contained at least one co-expression of interacting protein pairs in the RAD17–RFC-9-1-1 module with Pearson’s *r* values of ≥0.5. Figure 7G shows that the distribution of CEs of 16 PPIs in the RAD17–RFC-9-1-1 module for these 309 sets. CEs of 10 core PPIs were significantly higher than those of the three ring PPIs. For example, CE of RFC2 and RFC5 was 0.74, with Pearson’s *r* values of ≥0.5 in 229 gene expression sets among 309 sets. The ring proteins RAD9A, RAD1, and HUS1 of the RAD17–RFC-9-1-1 module form a PCNA-like ring structure that may interact with RFC-like complexes to regulate DNA binding in ATP-dependent or ATP-independent manners (Bermudez et al. 2003).

Figure 8 shows the module dynamics of the RAD17–RFC-9-1-1 supermodule during DNA replication from 309 gene expression sets, which are recorded in the GEO database and include all 8 proteins of this supermodule. Based on these 309 gene expression sets, we inferred seven modules that were described in more than three gene expression sets. For example, the RAF2-5 module (module 1) includes RCF2, RCF3, RCF4, and RCF5, and co-expressions of six PPIs (e.g., RCF2–RCF3, RCF2–RCF4, and RCF2–RCF5) with Pearson’s *r* values of ≥0.5 in 17 gene expression sets. Conversely, Pearson’s *r* values of the other 10 PPIs (e.g., RAD17–RFC2 and HUS1–RAD9A) of the RAD17–RFC-9-1-1 supermodule in these 17 sets were less than 0.5. Among these seven inferred modules, the RFC2-5 module (module 1, CORUM ID: 2200) and the RAD17–RFC-9-1-1 module (module 6, CORUM ID: 274) were recorded in the CORUM database and were derived from 17 and 5 gene expression sets, respectively. Inferred module 5, namely the RAF2–RAF4–RAF5 module, has been studied for DNA-dependent ATPase activity stimulated by PCNA (similar to the five-subunit RFC) and can unload PCNA from singly nicked circular DNA (Cai et al. 1997). In addition, we found that the RAD17–RFC module included the five proteins (i.e., RAD17 and RFC2-5), and interacts with RAD1 to form module 3, with RAD9A to form module 4, with RAD1 and HUS1 to form module 2, and with RAD9A and RAD1 to form module 7 for the regulation of DNA damage checkpoint response (Bermudez et al. 2003). Interestingly, according to these 309 sets, the RAD17–RFC module did not interact with HUS1 to form a module, and this was in agreement with a previous study (Bermudez et al. 2003). During DNA replication, the RFC2-5 module (module 1) and the RFC2–RFC4–RFC5 module (module 5) possess DNA-dependent ATPase activity and are not responsive to the addition of PCNA (Ellison and Stillman 1998) (Fig. 8). In the early steps of DNA damage recognition, the RAD17–RFC module (CORUM ID: 270) activates the checkpoint response (Lindsey-Boltz et al. 2001), and then binds to nicked circular, gapped, and primed DNA to recruit the RAD9A–RAD1–HUS1 module (module 6; CORUM ID: 274) for ATP-dependent DNA damage sensor (Bermudez et al. 2003). These results indicate that RFC2, RFC3, RFC4, and RFC5 play major roles in DNA damage recognition and that the RAD9, RAD1, and HUS1 could regulate them to bind to DNA with or without ATP. Interestingly, the core protein RAD17 forms the bridge between core and ring components, and co-expressions of the three core PPIs (i.e., RFC2-RAD17, RFC3-RAD17 and RFC4-RAD17) are slightly lower than those of the other core PPIs.

**Figure 8.**
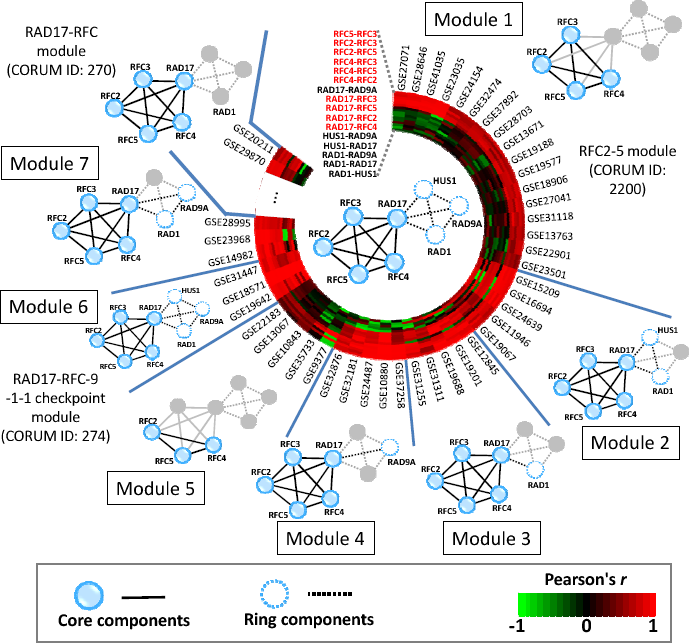
Module dynamics of the RAD17–RFC-9-1-1 supermodule during DNA replication. Among 309 gene expression sets, the RAD17–RFC-9-1-1 supermodule comprised seven modules with ≥3 gene expression sets containing all genes for RFC2–5, RFC7, RAD1, RAD9A, and HUS1. For example, modules 1 and 2 were presented in 17 and 5 sets, respectively. Co-expressions of 16 PPIs were measured using Pearson’s correlation *r* values ranging from −1 (green) to 1 (red). Proteins and PPIs with low gene co-expression (Pearson’s *r* < 0.5) are shown as gray circles and gray lines. Other proteins and PPIs are indicated with blue circles and black lines. For example, for module 5, Pearson’s *r* values of the three PPIs RFC2–RFC4, RFC2–RFC, and RFC4–RFC5 are more than 0.5 (red). Among these seven modules, only two (modules 1 and 6) are recorded in the CORUM database. The module dynamics provide cues for DNA damage sensors during DNA replication in an ATP-dependent manner. Solid circles/lines denote core proteins/PPIs and dashed circles/lines denote ring proteins/PPIs.

### Conclusions

In this work, we propose PPI evolution score and interface evolution score to elucidate module organizational principles and protein/module dynamics in biological networks using homologous modules and PPIs from complete genomic database. We identified core and ring components of modules, and showed similar prevalence of these in modules, despite differing module sizes. Compared with ring proteins and PPIs, core proteins and PPIs are often conserved in multiple species and taxonomic divisions, and participate in temporal and spatial cellular functions of most modules of supermodules. From biological network views, core components were shown to form hubs of biological networks, with high degrees and key roles in biological functions. Core proteins were often essential elements for survival. Conversely, ring proteins and PPIs were less conserved than core components and were involved in the biological functions of few modules within supermodules. From spatio-temporal view, core components were more frequently co-expressed and dynamic within modules than ring components according to 7,208 human gene expression sets. Based on these protein/module dynamics and gene co-expressions, we showed that core components are essential for biological functions of a module, whereas ring components contributed to the functions of a module in some cases. In summary, we propose the module organizational principles as follows: 1) a module comprises core and ring components that are conserved and non-conserved in multiple species during evolution, respectively; 2) core components often play major roles in the biological functions of modules, whereas ring components collaborate with core components to perform certain functions in only some cases; and 3) core components are more essential to temporal and spatial module dynamics and functions than ring components. The present definitions of core and ring components of modules are indicative of module organizational principles during evolution, and reflect cellular functions and network topology.

## Methods

### Homologous modules

Here, we used the module template M (including proteins A, B, C, and D) with six interfaces A–B, A–C, A–D, B–C, B–D, and C–D as an example (Fig. 1), and the homologous module of M was defined as follows: 1) A′, B′, C′, and D′ are homologous proteins of A, B, C, and D, respectively, with statistically significant sequence similarities (BLASTP *E*-values ≤ 10^−10^) (Matthews et al. 2001; Yu et al. 2004); 2) A′–B′, A′–C′, A′–D′, B′–C′, B′–D′, and C′–D′ are the best-matching homologous PPIs of A–B, A–C, A–D, B–C, B–D, and C–D, respectively, with statistically significant joint sequence similarities (joint *E*-value ≤ 10^−40^) (Chen et al. 2009); 3) A′, B′, C′, and D′ is the homologous module of template M, as indicated by high topological similarity (protein-aligned ratio of ≥0.5 and PPI-aligned ratio of ≥0.3). Protein- and PPI-aligned ratios were defined as the number of proteins or PPIs in the homologous module divided by the number of proteins or PPIs in the module template, respectively. Protein-aligned ratios of ≥0.5 and PPI-aligned ratios of ≥0.3 indicated topological similarity according to statistical analyses of 75,706 modules (370 reference modules) in 1,442 species based on the KEGG MODULE database (Kanehisa et al. 2008) (Supplemental Fig. S1).

### PPI evolution score and protein interface evolution score

We propose the PPI evolution score (PPIES) and protein interface evolution score (IES) to identify core and ring components of a module. To compute the PPIES of a PPI in a module family, we clustered National Center for Biotechnology Information (NCBI) taxonomy (Sayers et al. 2011) into six taxonomic divisions: mammals (MAM), vertebrates (VRT), invertebrates (INV), plants (PLN), bacteria (BCT), and archaea (ARC) (Supplemental Table S2). For each PPI *z* of a module family, PPIES was defined as

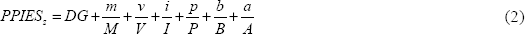

where *DG* is the number of taxonomic divisions that contain at least one species in homologous PPIs of the PPI *z* (Fig. 1D); *M*, *V*, *I*, *P*, *B*, and *A* are the total numbers of species of homologous modules belonging to MAM, VRT, INV, PLN, BCT, and ARC, in the module family, and *m*, *v*, *i*, *p*, *b*, and *a* are the numbers of species belonging to their respective taxonomic divisions of homologous PPIs of the PPI *z*, respectively (Fig. 1E). For each protein *k* in a module family, IES was set to the maximum PPIES, and was defined as *IES_k_* = max_1≤*j*≤*g*_(*PPIES_j_*), where *g* is the number of proteins that interact with protein *k*. Here, we considered proteins with IES ≥7 and PPIs with PPIES ≥7 as core components of a module; and all other proteins and PPIs were considered ring components. To evaluate conservation of modules during evolution for each module *d* in a module family, module evolution score (MES) was set to the mean PPIES and was defined as 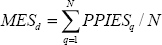, where *N* is the number of PPIs within module *d*.

### Protein-protein interactions in gene expression profiles

Proteins and PPIs change over time to assemble and disassemble a module for executing biological processes. Here, we quantified the dynamics of proteins and PPIs in time and space by assessing correlations between expression profiles of interacting proteins in 7,208 gene expression data sets (≥3 samples) derived from GEO (Barrett et al. 2009) (Fig. 6). To avoid the influence of genes with low expression and variance, we selected the gene *j* in a gene expression set based on the following criteria: average expression (*E*̄_j_) ≥ to the mean expression of all genes 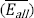 in a gene expression set; or the standard deviation of expression (S*_j_*) ≥ to the standard deviation of expression values for all genes (S_*all*_) in the gene expression set. For each module, we collected expression profiles contained expression values of all proteins in this module, and then calculated Pearson’s *r* values for each PPI within the module to construct correlation matrix. Here, we assume that an active module performed biological functions in a cell if at least one PPI of the module had high Pearson’s r ≥ h (here, h was set at 0.3, 0.5, or 0.7). For a PPI *p* (proteins *i* and *j*) in an active module, the co-expression ratio (CE) at the threshold *h* is defined as 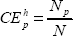, where *N* is the total number of these 7,208 expression profiles with at least one high co-expression (Pearson’s *r* ≥ *h*) of any PPI of this module; and *N_p_* is the number of expression profiles containing high co-expression of proteins *i* and *j* with Pearson’s *r* values of ≥*h*. For example, the CE of CDK1–CCNB1 is 0.76, reflecting high co-expression (Pearson’s *r* ≥ 0.5) in 825 of 1,085 gene expression sets when *h* = 0.5 (Fig. 1I).

## Acknowledgements

This paper was supported by National Science Council, partial supports of Ministry of Education and National Health Research Institutes (NHRI-EX100-10009PI). This paper is also particularly supported by "Aim for the Top University Plan" of the National Chiao Tung University and Ministry of Education. J.-M. Yang also thanks Core Facility for Protein Structural Analysis supported by National Core Facility Program for Biotechnology.

